# Exploring the impact of mitonuclear discordance on disease in Latin American admixed populations

**DOI:** 10.1101/2025.01.13.632776

**Authors:** Mauricio Ruiz, Daniela Böhme, Gabriela M. Repetto, Boris Rebolledo-Jaramillo

## Abstract

The coevolution of nuclear and mitochondrial genomes has guaranteed mitochondrial function for millions of years. The introduction of European (EUR) and African (AFR) genomes into the Ameridian continent during the Columbus exchange in Latin America created an opportunity to naturally test different combinations of nuclear and mitochondrial genomes. However, the impact of potential “mitonuclear discordance” (MND, differences in ancestries) has not been evaluated in Latin American admixed individuals (AMR) affected with developmental disorders, even though MND alters mitochondrial function and reduces viability in other organisms. To characterize MND in healthy and affected AMR individuals, we used AMR genotype data from the 1000 Genomes Project (n=385), two cohorts of 22q.11 deletion syndrome patients 22qDS-ARG (n=26) and 22qDS-CHL (n=58), and a cohort of patients with multiple congenital anomalies and/or neurodevelopmental disorders (DECIPHERD, n=172). Based on their importance to mitochondrial function, genes were divided into all mitonuclear genes (n=1035), high-mt (n=167), low-mt (n=793), or OXPHOS (n=169). We calculated local ancestry using FLARE, and estimated MND as the fraction of nuclear mitochondrial genes ancestry not matching the mtDNA ancestry, and ΔMND as (MND_offspring_ - MND_mother_)/MND_mother_. Generally, MND showed population and haplogroup distinctive distributions (ANOVA p<0.05), with haplogroup D showing the lowest MND 0.49±0.17 (mean ± s.d.). MND was significantly lower in 22qDS-ARG patients 0.43±0.24 and DECIPHERD patients 0.56±0.12 compared to healthy individuals 0.60±0.09 (ANOVA p<0.05). OXPHOS and high-mt showed the same trend, but with greater differences between healthy and affected individuals. MND seems to inform population history, and constraint among affected individuals, especially for OXPHOS and high-mt genes.

**Author summary:** We investigated the role of mitonuclear discordance (MND) in disease among Latin American admixed populations, describing how mismatches between nuclear and mitochondrial ancestries could impact mitochondrial function and disease phenotypes. Utilizing datasets from the 1000 Genomes Project and cohorts with rare genetic disorders, we demonstrated population- and haplogroup-specific patterns of MND, with haplogroup A showing the highest MND values. Affected individuals exhibited significantly lower MND than healthy controls, particularly in genes related to oxidative phosphorylation, suggesting potential links between MND and disease. Intergenerational analyses also revealed constraints on MND, pointing to selective pressures in disease contexts. These findings underscore the importance of mitonuclear interactions in shaping disease phenotypes and highlight the need for further research into their functional and clinical implications, particularly in underrepresented populations.

## Introduction

### Mitochondrial function depends on nuclear and mitochondrial factors

The mitochondrion (plural: mitochondria), is a double-layered organelle found in most eukaryotic cells. They are best known for their role in energy production, but they also participate in a variety of other cellular functions, including heme synthesis, calcium homeostasis, innate immunity, apoptosis, and the production of metabolic precursors for lipids, proteins, and nucleotides [1], [2], [3], [4], [5]. The mitochondrion is the only organelle in animal cells with its own genome: the mitochondrial DNA (mtDNA), a 16.5 kilobase (kb) circular double-stranded DNA molecule reflective of their past as prokaryotic organisms [6], [7]. In humans, inheritance of the mtDNA is non-Mendelian: the offspring receives the mtDNA exclusively from the mother. Mitochondrial proteins account for about 10% of the proteome [8]. The vast majority of mitochondrial proteins (∼1,000) are synthesized from nuclear genes and later transported to the mitochondrial matrix to perform their function [9], [10]. However, 13 essential subunits of respiratory complexes I, III, IV and V, 22 tRNAs and 2 rRNA of the mitochondrial specific translation machinery are synthesized within the mitochondria from mtDNA genes. Consequently, the assembly of the respiratory chain complexes requires the coordinated expression of nuclear and mitochondrial factors, and a tight regulation that has been maintained by mitonuclear coevolution over time [11].

### Mitonuclear discordance leads to dysfunctional mitochondria

Mitonuclear coevolution has been demonstrated using controlled crosses of non-human model organisms. Studies in mice [12], [13], fruit flies [14], and yeast [15] have demonstrated that mismatch between the nuclear and mitochondrial ancestries, from now on referred as mitonuclear discordance or MND, can cause mitonuclear incompatibility. This incompatibility is reflected by reduced viability and fecundity of interpopulation hybrids due to altered OXPHOS function [12], [13], [14], [15]. Fitness is usually restored if the hybrids are backcrossed with the maternal line, but not with the paternal line, supporting the hypothesis that their reduced fitness was caused by mitonuclear discordance [16]. Mitonuclear coevolution has also been observed in naturally occurring populations. In the eastern yellow robin, a type of bird, and killifish, a type of fish, it is believed that mitonuclear incompatibility has shaped their species differentiation [17], [18]. In humans, it has been postulated that mitonuclear incompatibility could occur due to the admixture of previously isolated populations, but its effect has not been properly studied [19], [20].

### Admixture leads to mitonuclear discordance in humans

The high mutation rate of the mtDNA [21] has resulted in the sequential accumulation of mtDNA variants (mtDNA haplogroups), reflective of human migration over time. The human mtDNA tree is rooted in Africa about 130-170,000 years before present (YBP). For the first 100,000 years, the mtDNA diversified within Africa giving rise to African-specific mtDNA haplogroups that are grouped into the macrohaplogroup L. Around 45,000 and 65,000 YBP, the macrohaplogroups M and N emerged and left Africa to successfully populate primarily Europe and Asia. From them, haplogroups A, B, C and D originated and only they populated the Americas about 20,000 YBP, suggesting that there were selective forces, possibly modulated by environmental adaptations, to constrain the radiation of other haplogroups throughout the Americas [22]. Genetic diversification due to physical isolation has been the dominant mode of human evolution since the emergence from Africa [23]. This gave rise to distinct continental groups: African, Asian, European and American, but upon arrival of Columbus to the American continent, a mere 532 years ago, the African, European, and Native American populations came into close and sustained contact for the first time [24]. Consequently, these diverse populations began to mix, creating novel combinations of nuclear and mitochondrial genotypes that had previously never existed together. Today, we find these novel combinations present in modern Latin-American admixed individuals [23], [25], but the potential contribution to health and disease of these admixtures has not been thoroughly considered.

### The effect of mitonuclear discordance due to admixture is unclear

The development of mitochondrial replacement therapy (MRT), an assisted reproductive technology, raised questions about the possibility of mitonuclear incompatibility in humans due to the artificial mixing of nuclear and mitochondrial genotypes [19]. In one of the strategies for MRT, the biparental nuclear genome is removed from a fertilized oocyte with diseased mitochondria and injected into an enucleated donor egg that contains healthy mitochondria. The process effectively generates a “three-parents” baby, due to the genetic contributions from two mothers and one father [19], [26]. Rishishwar and Jordan (2017) [27] addressed the concerns about potential mitonuclear incompatibilities introduced by MRT. They described the extent of naturally occurring MND in the 1000 Genomes Project Data, a cohort of 2504 presumably healthy individuals representing 26 populations across five continental groups [28]. They calculated the pairwise genetic distance between the individuals, and compared the differentiation obtained using nuclear and mitochondrial genotypes. As expected, they found evidence of mitonuclear coevolution, but they also observed multiples examples of coexistence of nuclear and mitochondrial genomes from divergent populations (admixture) and concluded that MND is not likely to cause mitonuclear incompatibilities or jeopardize the safety of MRT [19]. However, another study using a different research approach to understand MND in the same group of individuals found contrasting results. Zaidi and Makova (2019) used global and local ancestry inference to calculate MND as the proportion of nuclear ancestry not explained by the mtDNA ancestry. They observed that increased MND negatively correlated with mtDNA copy number (CN) [20]. mtDNA CN has been proposed as a biomarker for Parkinson’s disease [29], diabetes [30], and cancer [31]. Whether this correlation with MND can be extrapolated to understand these diseases needs further research, but this correlation demonstrates that MND can affect mitochondrial homeostasis. The main drawback of Rishishwar and Jordan (2017) and Zaidi and Makova (2019) articles is that the assessment of unrelated healthy adults provides no insights into whether selection had eliminated mitonuclear incompatibilities at earlier life stages [32]. This problem has been addressed in a large population study using mother-offspring pairs from the Genomics England 100,000 Genomes Project data. Wei and colleagues (2019)[33] compared groups of people of European and Asian descent with matching and unmatching nuclear and mitochondrial ancestries. They fitted a regression model to predict the transmission of European and Asian specific mtDNA haplogroup variants based on the ancestry of the nuclear genome. The results indicated that variants in the mismatched group were significantly more likely to match the ancestry of the nuclear genome than the ancestry of the mtDNA in which the variant occurred, suggesting, although indirectly, that there are mechanisms actively shaping the consistency of nuclear and mitochondrial ancestries in the maternal germline [34]. To date, these are the most exhaustive analyses of MND in humans, yet they provide inconclusive information on consequences.

### Mitonuclear interactions can modulate disease phenotypes

Traditionally, mtDNA variation has been ignored in GWAS. Only just recently nuclear and mitochondrial variants have been integrated to understand the contribution of mitonuclear interactions to disease risk [35]. This is illustrated by studies in neurodegenerative disorders, such as Parkinson’s and Alzheimer’s disease, where the inclusion of mtDNA haplogroups improved the predictions of polygenic risk scores built with nuclear mitochondrial variants [36], [37]; or a study evaluating the disruption of the physical interaction between nuclear and mitochondrial proteins of OXPHOS complex I, due to common genetic variation associated with risk of type 2 diabetes mellitus in Ashkenazi Jews [38].

Genotypes and phenotypes do not always have a linear relationship; thus, the identification of the genetic cause of a disease is not always straightforward. Even for rare Mendelian disorders with a known cause, the disease can display variable phenotypes among unrelated patients as well as among members of the same family. This variability can be partially explained by unknown genetic interactions contributed by so called “genetic modifiers”: genetic variants that can modify the phenotypic outcome of the primary disease-causing variant. The degree of the modifier effect can vary, which may result in large phenotypic variability of a disease [39].

### 22q11.2 deletion syndrome as an example of mitonuclear interactions

An example of this phenotypic variability can be found with the 22q11.2 deletion syndrome (22q11.2DS), a rare disorder with an estimated incidence of 1 in 4,000 live births [40]. The 22q11.2 deletion usually occurs *de novo* but is found to be inherited in about 10% of the cases. Phenotypes include congenital heart disease, palatal anomalies, immune compromise, developmental delay and learning difficulties, among many other characteristics that vary patient to patient [41], and even among members of the same family [42], [43]. Due to the 22q11.2 deletion, a copy of six nuclear mitochondrial genes is lost [40], [44], suggesting the involvement of mitonuclear interactions in the etiology of the disease. In fact, it has been observed that haploinsufficiency of *MRLP40*, one of the six nuclear encoded mitochondrial genes, impacts mitochondrial function in iPSC-derived neurons from 22q11.2DS patients [45]. In cases of maternally transmitted 22q11.2 deletion, since the offspring inherits only the paternal copies of the six nuclear encoded mitochondrial genes, we hypothesize that increased mitonuclear discordance contributes to the variable expression of the disease phenotypes in 22q11.2DS patients, and more generally, in affected Latin-American admixed patients. Thus, we describe mitonuclear discordance in admixed patients and compare it between healthy and disease cohorts to gauge the contribution on MND to disease.

## Methodology

### Samples

We reused data from multiple sequencing projects. We analyzed the genome of 385 samples of Latin-American admixed ancestry from the 1000 Genome Project Phase 3 dataset (1kGP-AMR) [46], which included 98 mother-offspring pairs (1kGP-pairs); the exome of 172 Chilean samples from the Decoding Complex Inherited Phenotypes in Rare Disorders cohort (DECIPHERD, referred as DRD for short) [47], a cohort where probands are affected by unknown rare disorders, causing congenital anomalies and/or neurodevelopmental disorders, which included 75 proband-only (DRD-affected) and 33 trios, from which healthy parents were used as controls (DRD-healthy), and mothers with offspring as pairs (DRD-pairs); the exome of 58 Chilean 22q11.2 deletion syndrome (22qDS) probands (22q-CHL); and the exome of 26 Argentinian 22qDS patients (22q-ARG), which included 12 mother-offspring pairs (22q-ARG-pairs) [48] (see Table 1).

**Table 1.** Datasets used for the calculation of baseline and disease-associated MND in LatinAmerican individuals.

| Dataset origin | Short name | # of samples | # pairs/trios | Usage |
| --- | --- | --- | --- | --- |
| 1000 Genomes Project Phase 3 | 1kGP-ALL | 3202 | 602 | To determine AMR, EUR and AFR haplotype references |
| 1000 Genomes Project Phase 3 | 1kGP-AMR | 385 | 98 | To describe MND and ΔMND in healthy LatinAmerican individuals |
| DECIPHERD | DRD-healthy | 63 | 0 | To describe MND in healthy Chilean individuals |
| DECIPHERD | DRD-affected | 75 | 0 | To describe MND in affected Chilean individuals |
| DECIPHERD | DRD-pairs | 66 | 33 | To describe ΔMND in affected Chilean mother-offspring pairs |
| 22q11.2DS - Chile | 22q-CHL | 58 | 0 | To describe MND in a known disease |
| 22q11.2DS - Argentina | 22q-ARG | 26 | 12 | To describe MND and ΔMND in a known disease |

### Ancestry inference

We calculated global ancestry with ADMIXTURE v.1.3 [49] using k = 5, to determine which 1kGP samples (out of 3,202 samples) would serve as Native American (AMR), European (EUR) and African (AFR) population references (admixture ≤ 1% for EUR and AFR, ≤ 10% for AMR). The 1kGP dataset included 63,993,320 single nucleotides variants (SNV), but we focused on 14,804,207 SNVs known to inform Latin American admixed ancestry according to gnomAD v.3 [50]. Exome data was imputed with the Michigan Imputation Server 2 [51] and filtered using rsq = 0.3.Then, for each dataset, we calculated local ancestry using FLARE v.0.5.1 [52] with default parameters. Mitochondrial DNA haplogroups were calculated using the web version of Haplogrep [53] on the off-target mtDNA reads obtained from exome sequencing.

### Mitonuclear discordance

Mitonuclear discordance (MND) has been defined as the fraction of nuclear ancestry not matching the mtDNA ancestry [20]. For example, in an individual with global nuclear components 55% European, 40% Native American and 5% African, with Native American mtDNA haplogroup B, MND would be 60% (the sum of European and African components). For this work, we calculated MND for each gene independently. Since each gene was represented by a set of SNPs of known local ancestry, we calculated the proportion of SNPs not matching the haplogroup’s ancestry and called that the “gene-wise MND”. We then averaged the gene-wise MND over all genes belonging to a particular gene set, e.g. “all mitonuclear genes”, or “OXPHOS genes”, and compared different sample sets. By default, we show the results for all mitonuclear genes, unless otherwise noted. We also defined the intergenerational change in mitonuclear discordance as: ΔMND = (MND_offspring_ -MND_mother_)/MND_mother_.

### Gene sets

Nuclear-encoded mitochondrial genes were classified into “all mitonuclear” (n = 1035), “OXPHOS” (n = 169), according to MitoCarta v.3.0 [54], and “high-mt” (n = 167) or “low-mt” (n = 793), according to Sloan, et al. (2015) [55].

## Statistics

Results are shown as mean ± standard deviation, unless otherwise noted. Multiple group comparisons were calculated using ANOVA for all cases where normality or the central limit theorem were applicable, otherwise, we used Kruskal-Wallis. Pairwise comparisons were calculated with the t-test or Mann-Whitney U test, accordingly. Multiple hypotheses testing was controlled with Bonferroni’s correction. For all tests, the significance level alpha was set at 0.05. We used R v.4.3.3 to perform all statistical analyses and plots.

## Results

### Samples

We leveraged existing datasets to describe MND and ΔMND in healthy and affected individuals. Out of the 3,202 1000 Genomes Project Phase 3 individuals [46], 336 African, 198 European and 25 American individuals served as population references for all calculations. After imputation of exome data, we obtained 6,279,408 22q-ARG SNVs; 10,642,230 22q-CHL SNVs; 2,361,135 DRD-pairs and DRD-healthy SNVs; and 8,504,595 DRD-affected SNVs. We categorized the cohorts according to Table 1.

### MND in healthy individuals show strong population and haplogroup-specific distributions

First, we calculated mean MND distributions for all populations and observed strong population specificity, Puerto Rican (PUR, 0.75 ± 0.22); Colombian (CLM, 0.72 ± 0.16); Peruvian (PEL, 0.27 ± 0.18); Mexican (MXL, 0.49 ± 0.18); Chilean, (CHL 0.60 ± 0.09); ANOVA F = 112.7, p = 3.37E-66 (all pairwise comparisons were statistically significant, figure 1A), and the mean of the distributions positively correlated with European ancestry, rho = 0.689, p = 2.42E-64, S1 Fig. Then, we merged all Latin American haplogroups belonging to the same branch into single letter macrohaplogroups to increase sample size. We observed that macrohaplogroup A had the highest mean MND (0.67 ± 0.21), significantly different compared to macrohaplogroups B (0.55 ± 0.22), C (0.58 ± 0.22), and D (0.49 ± 0.17), which in turn had similar distributions, ANOVA F = 7.529 p = 7.7E-05. Noticeably, macrohaplogroup D had the lowest mean MND and variance. Pairwise comparisons are shown in figure 1B.

**Figure 1.**
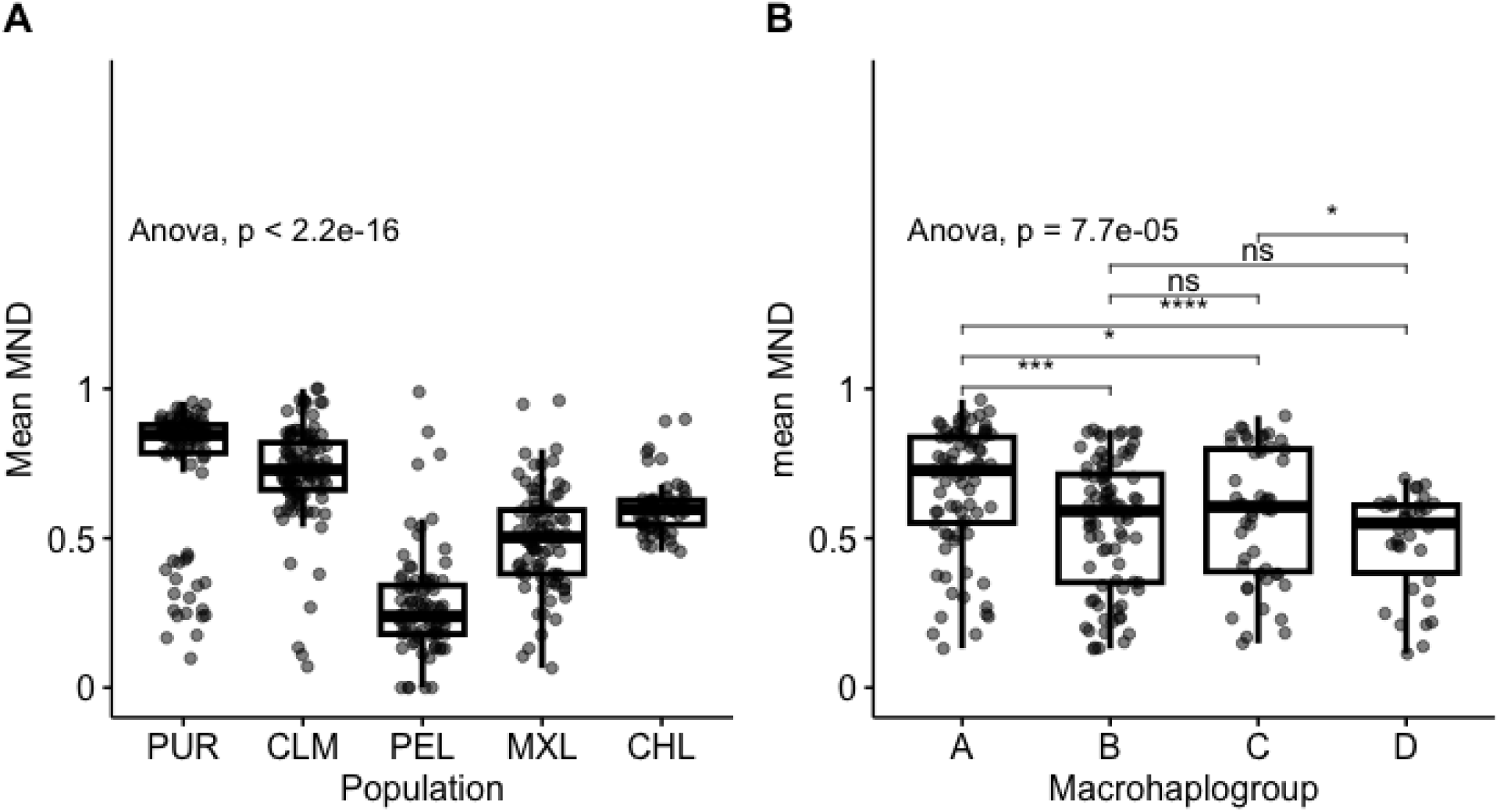
Population and haplogroup-specific MDN distribution. (A) Distribution of mean MND among 1kGP and Chilean populations. (B) Distribution of mean MND among Latin American macrohaplogroups.

### Affected individuals have lower MND

We compared the mean MND of healthy parents from the DECIPHERD cohort to the mean MND of affected cohorts. We observed that in general, using all mitonuclear genes, MND in affected individuals was lower: DRD-healthy (0.60 ± 0.09); DRD-affected (0.56 ± 0.12); 22q-CHL (0.57 ± 0.13), with the 22q-ARG cohort showing the lowest mean MND (0.43 ± 0.24), ANOVA F = 10.7, p = 1.4E-6. Interestingly, the variance of 22q-ARG patients was 3.6 times greater than the variance of their Chilean counterpart (0.058 and 0.016, respectively, p = 6.33E-5) (figure 2A). Similarly, we ran the same analyses, but focusing only on OXPHOS genes. We observed an even stronger difference between healthy and affected cohorts: healthy (0.62 ± 0.1), DRD-affected (0.55 ± 0.13), 22q-CHL (0.57 ± 0.13), and 22q-ARG (0.43 ± 0.25), ANOVA F = 11.6, p = 4.2E-7 (figure 2 B and Table 2). The results for high-mt and low-mt genes are shown in S2 Fig.

**Figure 2.**
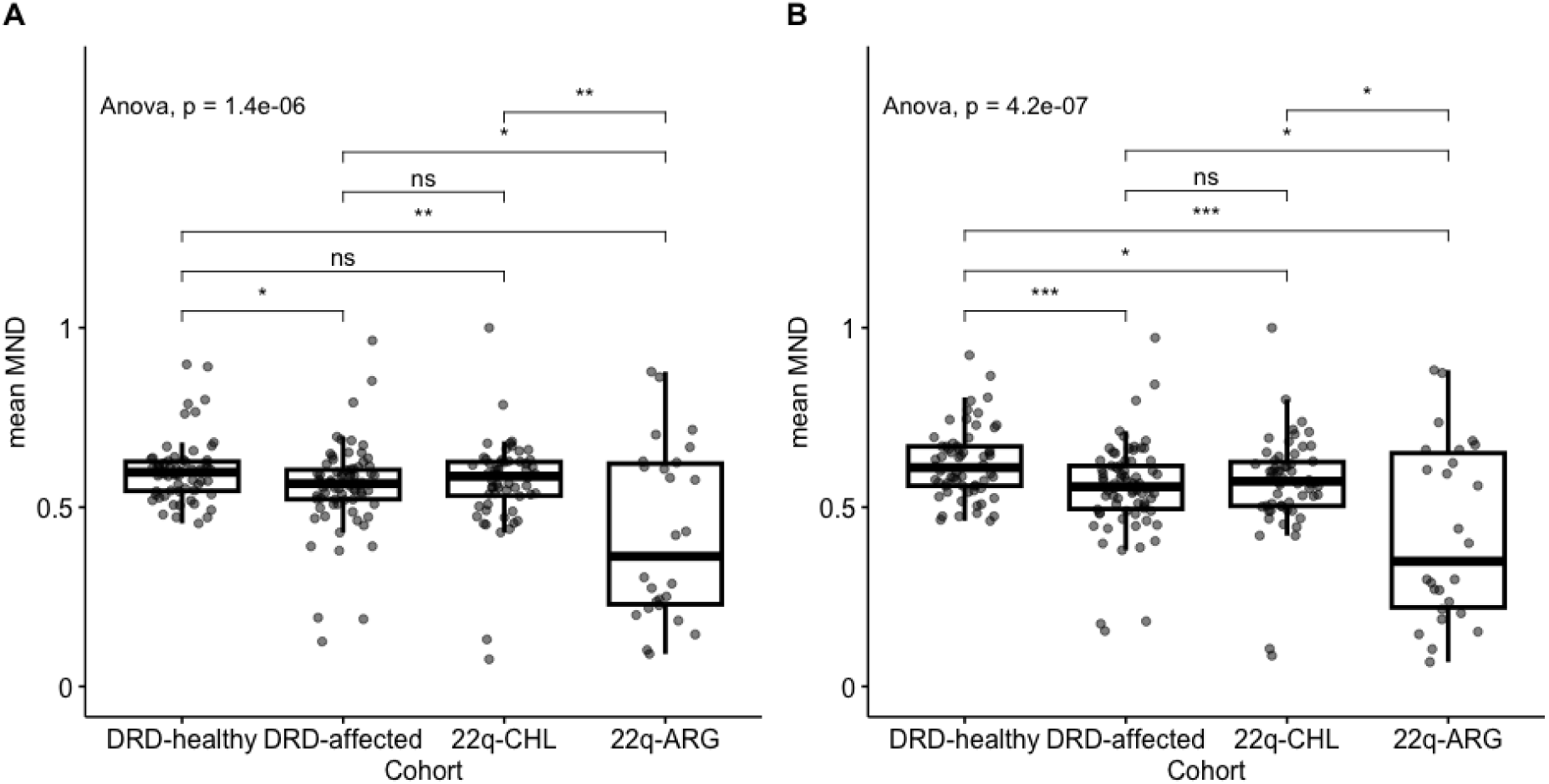
Mean MND in cohorts of patients with different genetic disorders. We compared healthy individuals to multiple cohorts of patients. The first three distribution correspond to Chilean individuals, whereas the last distribution correspond to patients from Argentina. (A) Comparison of mean MDN using all mitonuclear genes. (B) Comparison of mean MDN using OXPHOS genes.

**Table 2.**
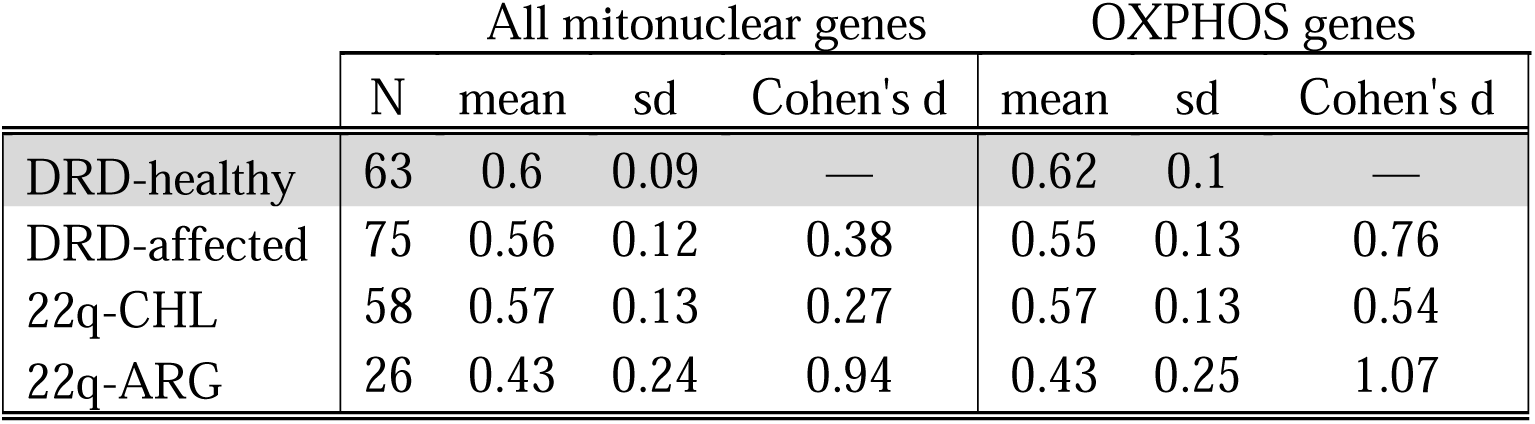
Greater effect sizes using OXPHOS genes. Effect sizes were calculated by comparing DRD-healthy to the other cohorts.

### Signs of constraint of ΔMND

We observed that the intergenerational change in MND had lower median in DRD -0.01 [-0.18 – 0.24] (median and range) and 22q-ARG pairs -0.05 [-0.43 – 0.50], compared to healthy 0.02 [-1.0 – 1.5], yet the distributions were not significantly different, Kruskal-Wallis p = 0.92. However, we observed significant differences in variance between healthy (s^2^ = 0.16) and DRD (s^2^ = 0.01) pairs, p = 7.0E-12, but not between healthy and 22q-ARG pairs (s^2^ = 0.06, p = 0.09), Figure 3.

**Figure 3.**
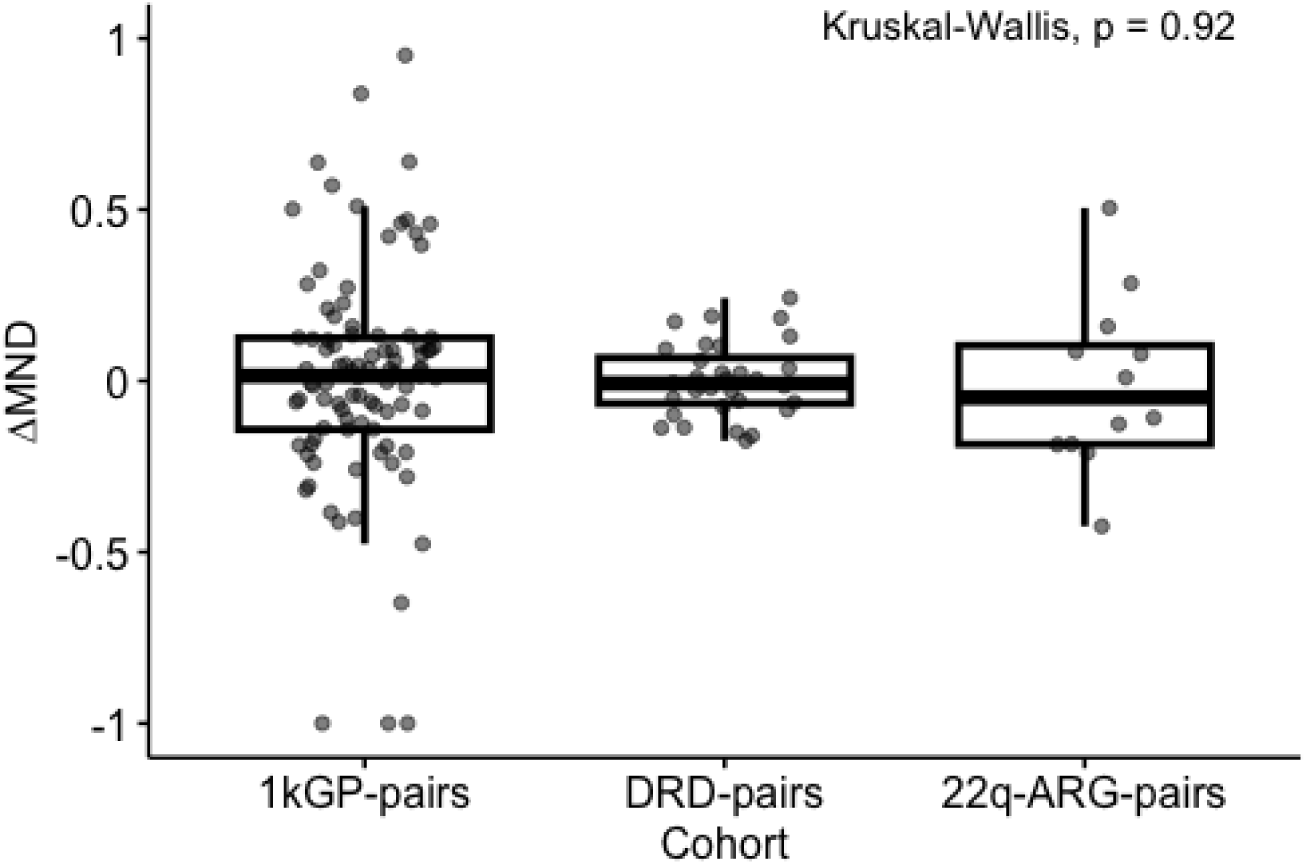
Comparison of ΔMND in affected cohorts. 1kGP-pairs represent healthy individuals, while DRD-pairs and 22q-ARG-pairs represent cohorts affected by rare genetic disorders.

## Discussion

This study explored the contribution of mitonuclear discordance to disease in Latin American admixed populations, offering a novel perspective on the interplay between nuclear and mitochondrial genomes in diverse human cohorts. By leveraging comprehensive datasets, including the 1000 Genomes Project and cohorts of patients with rare genetic disorders, this research highlights the potential impact of MND on mitochondrial function and disease phenotypes.

Our results showed strong population- and haplogroup-specific distributions of MND, with significant correlations between MND and European ancestry, results that align with previous studies. Notably, in our study, haplogroup D exhibited the lowest MND, suggesting potential evolutionary constraints. While direct studies on haplogroup D are limited, research has shown that certain mtDNA lineages may be subject to evolutionary constraints due to the interaction between haplogroup defining variants. For example, in a H7A haplogroup background, the J haplogroup defining variant m.13708G>A causes a complex phenotype including a combination of connective tissue, neurological, and metabolic symptoms, demonstrating that common nonpathogenic variation can cause mitochondrial dysfunction due to variants incompatibility [56], [57].

The observed lower mitonuclear discordance in affected individuals compared to healthy controls, particularly pronounced in OXPHOS genes, suggests that MND may influence disease phenotypes. This underscores the critical role of OXPHOS in maintaining mitochondrial homeostasis, suggesting that higher MND levels could be detrimental to survival. Research has demonstrated that mitochondrial dysfunction, including impaired OXPHOS, is implicated in various diseases. For instance, mitochondrial dysfunction has been linked to the etiology of bipolar disorder, with evidence pointing to dysregulated OXPHOS and altered brain bioenergetics [58]. Additionally, studies have shown that mutations affecting OXPHOS can lead to severe disorders. For example, combined oxidative phosphorylation deficiency type 14, caused by mutations in the *FARS2* gene, manifests in conditions such as epileptic status and other neurological impairments [59] – which could explain some of the neurological phenotypes observed in the DECIPHERD cohort. These findings highlight the importance of mitonuclear interactions in disease manifestation and the necessity of coordinated function between nuclear and mitochondrial genomes for optimal OXPHOS activity. Disruptions in this coordination, may compromise mitochondrial function and contribute to disease development.

We also examined intergenerational changes in MND, which revealed signs of constraint in affected cohorts. As expected, ΔMND in healthy individual showed signs of genetic drift, since the distribution was centered around zero. While the median ΔMND was lower in affected mother-offspring pairs, these differences were not stastically significant, yet differences in variance were noted. This difference in variance may reflect selective pressures or mechanisms that limit mitonuclear discordance across generations. Studies have shown that mitonuclear interactions are subject to selective pressures, particularly in the context of disease. For example, research on admixed human populations has demonstrated that mitonuclear DNA discordance can affect mitochondrial DNA copy number or gene expression, with higher discordance leading to lower mtDNA copy numbers and lower gene expression. This suggests that mitonuclear compatibility is crucial for maintaining mitochondrial function and that selective pressures may act to minimize discordance across generations [20], [60]. Additionally, studies on mitonuclear incompatibilities in allopatric speciation have highlighted that rapid evolution of the mitochondrial genome creates intrinsic selection pressures favoring nuclear gene mutations that maintain mitochondrial function. This indicates that mitonuclear interactions are under selective constraints to ensure compatibility, which may be particularly relevant in disease contexts where mitochondrial function is compromised [61].

Our results suggest that MND could act as a genetic modifier, contributing to phenotypic variability in diseases such as 22q11.2DS. This aligns with the growing recognition of mitonuclear interactions in modulating disease risk, especially in neurodegenerative, immunological, and developmental disorders. Our previous research demonstrated that mtDNA heteroplasmy may influence the incomplete penetrance of the palatal phenotype in 22q11.2DS, highlighting the potential role of mitochondrial variants within a specific genetic background as genetic modifiers [48]. Also, as mentioned earlier, research on admixed human populations has shown that mitonuclear DNA discordance correlates with variations in mtDNA gene expression levels, and mtDNA copy number, suggesting that mitonuclear interactions may act as genetic modifiers, influencing phenotypic variability in various diseases [60], [62]. Furthermore, research on Alzheimer’s disease has indicated that mitonuclear interactions influence disease risk. Associations between mitochondrial DNA haplogroups and nuclear-encoded mitochondrial genes have been linked to variations in dementia risk and age of onset, underscoring the significance of mitonuclear interactions in neurodegenerative diseases [36].

One of our study’s strengths lies in its use of diverse datasets, allowing for robust comparisons across populations and disease states, and the focus on Latin American cohorts which are traditionally underrepresented in genomic studies. While correlations were observed, mechanistic insights into how MND influences disease phenotypes require further investigation, particularly the exploration of functional consequences of MND in cellular and animal models, to hopefully develop clinical tools to assess MND as a potential biomarker for disease risk and progression.

## Acknowledgments

We would like to thank the patients and families who contributed their data to this research, as well as the team behind the DECIPHERD project and Hospital Garrahan.

## Supplementary information

**S1 Fig. Correlations between mean MND and European ancestry**

**S2 Fig. Mean MND in cohorts of patients with multiple genetic disorders**

## References

[1] A. J. Anderson, T. D. Jackson, D. A. Stroud, and D. Stojanovski, “Mitochondria—hubs for regulating cellular biochemistry: Emerging concepts and networks,” 2019. doi: 10.1098/rsob.190126.

[2] L. Craven, C. L. Alston, R. W. Taylor, and D. M. Turnbull, “Recent Advances in Mitochondrial Disease,” Annual Review of Genomics and Human Genetics, 2017, doi: 10.1146/annurev-genom-091416-035426.

[3] J. B. Spinelli and M. C. Haigis, “The multifaceted contributions of mitochondria to cellular metabolism,” 2018. doi: 10.1038/s41556-018-0124-1.

[4] D. C. Chan, “Mitochondrial Dynamics and Its Involvement in Disease,” Annual Review of Pathology: Mechanisms of Disease, 2020, doi: 10.1146/annurev-pathmechdis-012419-032711.

[5] C. S. Ahn and C. M. Metallo, “Mitochondria as biosynthetic factories for cancer proliferation,” 2015. doi: 10.1186/s40170-015-0128-2.

[6] I. Zachar and G. Boza, “Endosymbiosis before eukaryotes: mitochondrial establishment in protoeukaryotes,” 2020. doi: 10.1007/s00018-020-03462-6.

[7] L. Levin, A. Blumberg, G. Barshad, and D. Mishmar, “Mito-nuclear co-evolution: The positive and negative sides of functional ancient mutations,” Frontiers in Genetics, 2014, doi: 10.3389/fgene.2014.00448.

[8] S. Rath et al., “MitoCarta3.0: An updated mitochondrial proteome now with sub-organelle localization and pathway annotations,” Nucleic Acids Research, 2021, doi: 10.1093/nar/gkaa1011.

[9] J. B. Stewart and P. F. Chinnery, “Extreme heterogeneity of human mitochondrial DNA from organelles to populations,” 2020. doi: 10.1038/s41576-020-00284-x.

[10] J. Song, J. M. Herrmann, and T. Becker, “Quality control of the mitochondrial proteome,” 2021. doi: 10.1038/s41580-020-00300-2.

[11] M. T. Couvillion, I. C. Soto, G. Shipkovenska, and L. S. Churchman, “Synchronized mitochondrial and cytosolic translation programs,” Nature, 2016, doi: 10.1038/nature18015.

[12] H. Ma et al., “Incompatibility between Nuclear and Mitochondrial Genomes Contributes to an Interspecies Reproductive Barrier,” Cell Metabolism, 2016, doi: 10.1016/j.cmet.2016.06.012.

[13] A. Latorre-Pellicer et al., “Mitochondrial and nuclear DNA matching shapes metabolism and healthy ageing,” Nature, 2016, doi: 10.1038/nature18618.

[14] J. A. Mossman, J. G. Tross, N. A. Jourjine, N. Li, Z. Wu, and D. M. Rand, “Mitonuclear Interactions Mediate Transcriptional Responses to Hypoxia in Drosophila,” Molecular biology and evolution, 2017, doi: 10.1093/molbev/msw246.

[15] T. H. M. Nguyen, S. Sondhi, A. Ziesel, S. Paliwal, and H. L. Fiumera, “Mitochondrial-nuclear coadaptation revealed through mtDNA replacements in Saccharomyces cerevisiae,” BMC Evolutionary Biology, 2020, doi: 10.1186/s12862-020-01685-6.

[16] C. K. Ellison and R. S. Burton, “Interpopulation hybrid breakdown maps to the mitochondrial genome,” Evolution, 2008, doi: 10.1111/j.1558-5646.2007.00305.x.

[17] H. E. Morales et al., “Concordant divergence of mitogenomes and a mitonuclear gene cluster in bird lineages inhabiting different climates,” Nature Ecology and Evolution, 2018, doi: 10.1038/s41559-018-0606-3.

[18] T. Z. Baris et al., “Evolved genetic and phenotypic differences due to mitochondrial-nuclear interactions,” PLoS Genetics, 2017, doi: 10.1371/journal.pgen.1006517.

[19] L. Rishishwar and I. K. Jordan, “Implications of human evolution and admixture for mitochondrial replacement therapy,” BMC Genomics, 2017, doi: 10.1186/s12864-017-3539-3.

[20] A. A. Zaidi and K. D. Makova, “Investigating mitonuclear interactions in human admixed populations,” Nature Ecology and Evolution, 2019, doi: 10.1038/s41559-018-0766-1.

[21] B. Rebolledo-Jaramillo et al., “Maternal age effect and severe germ-line bottleneck in the inheritance of human mitochondrial DNA,” Proceedings of the National Academy of Sciences of the United States of America, 2014, doi: 10.1073/pnas.1409328111.

[22] D. C. Wallace, “Mitochondrial DNA Variation in Human Radiation and Disease,” 2015. doi: 10.1016/j.cell.2015.08.067.

[23] E. T. Norris et al., “Genetic ancestry, admixture and health determinants in Latin America,” BMC Genomics, 2018, doi: 10.1186/s12864-018-5195-7.

[24] I. K. Jordan, “The Columbian Exchange as a source of adaptive introgression in human populations,” Biology Direct, 2016, doi: 10.1186/s13062-016-0121-x.

[25] A. Ruiz-Linares et al., “Admixture in Latin America: Geographic Structure, Phenotypic Diversity and Self-Perception of Ancestry Based on 7,342 Individuals,” PLoS Genetics, 2014, doi: 10.1371/journal.pgen.1004572.

[26] A. Greenfield, P. Braude, F. Flinter, R. Lovell-Badge, C. Ogilvie, and A. C. F. Perry, “Assisted reproductive technologies to prevent human mitochondrial disease transmission,” 2017. doi: 10.1038/nbt.3997.

[27] L. Rishishwar and I. K. Jordan, “Implications of human evolution and admixture for mitochondrial replacement therapy,” BMC Genomics, 2017, doi: 10.1186/s12864-017-3539-3.

[28] A. Auton et al., “A global reference for human genetic variation,” 2015. doi: 10.1038/nature15393.

[29] A. Pyle, H. Anugrha, M. Kurzawa-Akanbi, A. Yarnall, D. Burn, and G. Hudson, “Reduced mitochondrial DNA copy number is a biomarker of Parkinson’s disease,” Neurobiology of Aging, 2015, doi: 10.1016/j.neurobiolaging.2015.10.033.

[30] H. K. Lee et al., “Decreased mitochondrial DNA content in peripheral blood precedes the development of non-insulin-dependent diabetes mellitus,” Diabetes Research and Clinical Practice, 1998, doi: 10.1016/S0168-8227(98)00110-7.

[31] M. Yu, “Generation, function and diagnostic value of mitochondrial DNA copy number alterations in human cancers,” 2011. doi: 10.1016/j.lfs.2011.05.010.

[32] G. E. Hill, J. C. Havird, D. B. Sloan, R. S. Burton, C. Greening, and D. K. Dowling, “Assessing the fitness consequences of mitonuclear interactions in natural populations,” Biological Reviews, 2019, doi: 10.1111/brv.12493.

[33] W. Wei et al., “Germline selection shapes human mitochondrial DNA diversity,” Science, 2019, doi: 10.1126/science.aau6520.

[34] W. Wei et al., “Germline selection shapes human mitochondrial DNA diversity,” Science, 2019, doi: 10.1126/science.aau6520.

[35] A. H. Ludwig-Słomczyńska et al., “Mitochondrial GWAS and association of nuclear - Mitochondrial epistasis with BMI in T1DM patients,” BMC Medical Genomics, 2020, doi: 10.1186/s12920-020-00752-7.

[36] S. J. Andrews et al., “Mitonuclear interactions influence Alzheimer’s disease risk,” Neurobiology of Aging, 2020, doi: 10.1016/j.neurobiolaging.2019.09.007.

[37] A. Schulmann et al., “Novel Complex Interactions between Mitochondrial and Nuclear DNA in Schizophrenia and Bipolar Disorder,” Molecular Neuropsychiatry, 2019, doi: 10.1159/000495658.

[38] M. Gershoni et al., “Disrupting mitochondrial-nuclear coevolution affects OXPHOS complex i integrity and impacts human health,” Genome Biology and Evolution, 2014, doi: 10.1093/gbe/evu208.

[39] K. M. T. H. Rahit and M. Tarailo-Graovac, “Genetic Modifiers and Rare Mendelian Disease,” 2020. doi: 10.3390/genes11030239.

[40] D. M. McDonald-McGinn et al., “22q11.2 deletion syndrome,” Nature Reviews Disease Primers, 2015, doi: 10.1038/nrdp.2015.71.

[41] R. D. Burnside, “22q11.21 deletion syndromes: A review of proximal, central, and distal deletions and their associated features,” 2015. doi: 10.1159/000438708.

[42] D. M. McDonald-McGinn et al., “Phenotype of the 22q11.2 deletion in individuals identified through an affected relative: Cast a wide FISHing net!,” in Genetics in Medicine, 2001. doi: 10.1097/00125817-200101000-00006.

[43] C. Cancrini et al., “Clinical features and follow-up in patients with 22q11.2 deletion syndrome,” Journal of Pediatrics, 2014, doi: 10.1016/j.jpeds.2014.01.056.

[44] P. Devaraju and S. S. Zakharenko, “Mitochondria in complex psychiatric disorders: Lessons from mouse models of 22q11.2 deletion syndrome,” BioEssays, 2017, doi: 10.1002/bies.201600177.

[45] J. Li et al., “Mitochondrial deficits in human iPSC-derived neurons from patients with 22q11.2 deletion syndrome and schizophrenia,” Translational Psychiatry, 2019, doi: 10.1038/s41398-019-0643-y.

[46] M. Byrska-Bishop et al., “High-coverage whole-genome sequencing of the expanded 1000 Genomes Project cohort including 602 trios,” Cell, vol. 185, no. 18, pp. 3426–3440.e19, Sep. 2022, doi: 10.1016/j.cell.2022.08.004.

[47] M. C. Poli et al., “Decoding complex inherited phenotypes in rare disorders: the DECIPHERD initiative for rare undiagnosed diseases in Chile,” Eur J Hum Genet, vol. 32, no. 10, pp. 1227–1237, Oct. 2024, doi: 10.1038/s41431-023-01523-5.

[48] B. Rebolledo-Jaramillo, M. G. Obregon, V. Huckstadt, A. Gomez, and G. M. Repetto, “Contribution of mitochondrial dna heteroplasmy to the congenital cardiac and palatal phenotypic variability in maternally transmitted 22q11.2 deletion syndrome,” Genes, 2021, doi: 10.3390/genes12010092.

[49] D. H. Alexander, J. Novembre, and K. Lange, “Fast model-based estimation of ancestry in unrelated individuals,” Genome Research, 2009, doi: 10.1101/gr.094052.109.

[50] S. Chen et al., “A genomic mutational constraint map using variation in 76,156 human genomes,” Nature, vol. 625, no. 7993, pp. 92–100, Jan. 2024, doi: 10.1038/s41586-023-06045-0.

[51] S. Das et al., “Next-generation genotype imputation service and methods,” Nat Genet, vol. 48, no. 10, pp. 1284–1287, Oct. 2016, doi: 10.1038/ng.3656.

[52] S. R. Browning, R. K. Waples, and B. L. Browning, “Fast, accurate local ancestry inference with FLARE,” The American Journal of Human Genetics, vol. 110, no. 2, pp. 326–335, Feb. 2023, doi: 10.1016/j.ajhg.2022.12.010.

[53] H. Weissensteiner et al., “HaploGrep 2: mitochondrial haplogroup classification in the era of high-throughput sequencing,” Nucleic acids research, 2016, doi: 10.1093/nar/gkw233.

[54] S. Rath et al., “MitoCarta3.0: An updated mitochondrial proteome now with sub-organelle localization and pathway annotations,” Nucleic Acids Research, 2021, doi: 10.1093/nar/gkaa1011.

[55] D. B. Sloan, P. D. Fields, and J. C. Havird, “Mitonuclear linkage disequilibrium in human populations”, doi: 10.1098/rspb.2015.1704.

[56] N. J. Lake et al., “Quantifying constraint in the human mitochondrial genome,” Nature, vol. 635, no. 8038, pp. 390–397, Nov. 2024, doi: 10.1038/s41586-024-08048-x.

[57] P. M. Schaefer et al., “Combination of common mtDNA variants results in mitochondrial dysfunction and a connective tissue dysregulation,” Proc. Natl. Acad. Sci. U.S.A., vol. 119, no. 45, p. e2212417119, Nov. 2022, doi: 10.1073/pnas.2212417119.

[58] S. Gonzalez, “The Role of Mitonuclear Incompatibility in Bipolar Disorder Susceptibility and Resilience Against Environmental Stressors,” Front. Genet., vol. 12, p. 636294, Mar. 2021, doi: 10.3389/fgene.2021.636294.

[59] X. Zhang, F. Xiang, D. Li, F. Yang, S. Yu, and X. Wang, “Adult-onset combined oxidative phosphorylation deficiency type 14 manifests as epileptic status: a new phenotype and literature review,” BMC Neurol, vol. 24, no. 1, p. 15, Jan. 2024, doi: 10.1186/s12883-023-03480-4.

[60] E. Torres-Gonzalez and K. D. Makova, “Exploring the Effects of Mitonuclear Interactions on Mitochondrial DNA Gene Expression in Humans,” Front. Genet., vol. 13, p. 797129, Jun. 2022, doi: 10.3389/fgene.2022.797129.

[61] R. S. Burton, “The role of mitonuclear incompatibilities in allopatric speciation,” Cell. Mol. Life Sci., vol. 79, no. 2, p. 103, Feb. 2022, doi: 10.1007/s00018-021-04059-3.

[62] A. A. Zaidi and K. D. Makova, “Investigating mitonuclear interactions in human admixed populations,” Nature Ecology and Evolution, 2019, doi: 10.1038/s41559-018-0766-1.

