## Supplementary figures for "Exploring the impact of mitonuclear discordance on disease in Latin American admixed populations"

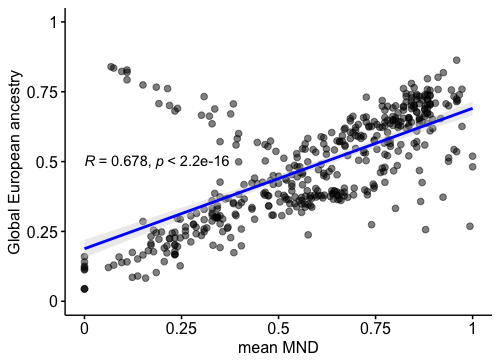

**Supp. Figure 1.** Scatterplot showing the correlation between mean MND and European ancestry. Linear regression line is shown in blue.

A	B

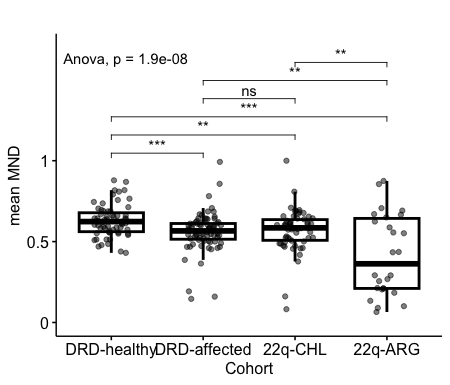

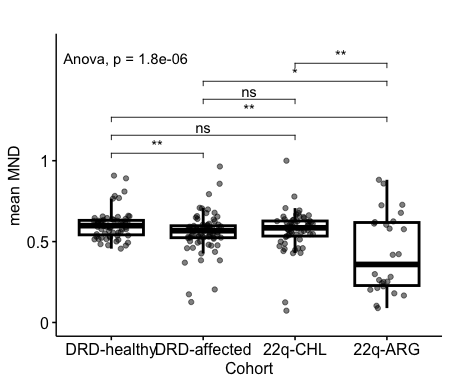

**Supp. Figure 2. Mean MND in cohorts of patients with different genetic disorders.** Similarly to figure 2 from the main text, we compared healthy individuals to multiple cohorts of patients. (A) Comparison of mean MDN using high-mt genes. (B) Comparison of mean MDN using low-mt genes.
